# SQANTI: extensive characterization of long read transcript sequences for quality control in full-length transcriptome identification and quantification

**DOI:** 10.1101/118083

**Authors:** Manuel Tardaguila, Lorena de la Fuente, Cristina Marti, Cécile Pereira, Francisco Jose Pardo-Palacios, Hector del Risco, Marc Ferrell, Maravillas Mellado, Marissa Macchietto, Kenneth Verheggen, Mariola Edelmann, Iakes Ezkurdia, Jesus Vazquez, Michael Tress, Ali Mortazavi, Lennart Martens, Susana Rodriguez-Navarro, Victoria Moreno, Ana Conesa

## Abstract

High-throughput sequencing of full-length transcripts using long reads has paved the way for the discovery of thousands of novel transcripts, even in very well annotated organisms as mice and humans. Nonetheless, there is a need for studies and tools that characterize these novel isoforms. Here we present SQANTI, an automated pipeline for the classification of long-read transcripts that computes 47 descriptors that can be used to assess the quality of the data and of the preprocessing pipelines. We applied SQANTI to a neuronal mouse transcriptome using PacBio long reads and illustrate how the tool is effective in readily describing the composition of and characterizing the full-length transcriptome. We perform extensive evaluation of ToFU PacBio transcripts by PCR to reveal that an important number of the novel transcripts are technical artifacts of the sequencing approach, and that SQANTI quality descriptors can be used to engineer a filtering strategy to remove them. Most novel transcripts in this curated transcriptome are novel combinations of existing splice sites, result more frequently in novel ORFs than novel UTRs and are enriched in both general metabolic and neural specific functions. We show that these new transcripts have a major impact in the correct quantification of transcript levels by state-of-the-art short-read based quantification algorithms. By comparing our iso-transcriptome with public proteomics databases we find that alternative isoforms are elusive to proteogenomics detection and are variable in protein changes with respect to the principal isoform of their genes. SQANTI allows the user to maximize the analytical outcome of long read technologies by providing the tools to deliver quality-evaluated and curated full-length transcriptomes. SQANTI is available at <u>https://bitbucket.org/ConesaLab/sqanti</u>.

## INTRODUCTION

Alternative Splicing (AS) and Alternative Polyadenylation (APA) are among the most fascinating and challenging aspects of eukaryotic transcriptomes. AS and APA are considered major mechanisms to generate transcriptome complexity and thus expand proteome diversity of higher organisms^1–3^. These post-transcriptional mechanisms have been reported to play critical roles in differentiation^4–7^, speciation^3,8^ and multiple human diseases such as cancer^9–11^, diabetes^12,13^ or neurological disorders^14–18^, and therefore play a fundamental role in the establishment of organismal complexity^3,19,20^. The genome-wide analysis of AS has been done primarily using first exon microarrays and more recently short-read RNA-seq. These two methods are effective for the identification of AS events such as exon skipping or intron retention and have established the involvement of AS in many biological processes. However, both technologies have serious limitations for the reconstruction of the actual expressed transcripts, as short reads break the continuity of the transcript sequences and fail to resolve assembly ambiguities at complex loci^21,22^. This impairs any studies that would catalogue specific transcriptomes, investigate cis-acting mechanisms within transcripts, infer open reading frames or understand functional aspects of isoform diversity.

There has been increasing interest in the application of single-molecule sequencing to obtain full-length transcripts in animals and plants as long reads eliminate the need for short-read assembly and allow direct isoform sequencing. Currently there exists three different long read transcriptome sequencing platforms; PacBio^22–24^, Moleculo^25^ and Nanopore^26^ Here, we have used the popular PacBio Iso-Seq protocol, which consists of full-length cDNA enrichment using the ClonTech SMARTer kit followed by building single molecule SMRTbell libraries with specific PacBio linkers that are subsequently sequenced. PacBio reads are typically longer than the full-length cDNA sequence, which means that each molecule can go through several passes of sequencing. The consensus of these passes is called a Read of Insert (RoI), which is the current standard PacBio output. RoIs where both cDNA primers and the poly(A) can be identified are called Full-length (FL) reads, while those that miss any of these tags are deemed non Full-length reads. PacBio sequencing suffers, however, from a relatively high raw error rate (∼15%^27^) and a lower throughput compared to Illumina. There are several described methods for PacBio error correction and transcript identification. Au et al^28^, proposed a hybrid sequencing approach (IDP), where PacBio RoIs are first corrected with the more accurate Illumina reads using the computationally intensive LSC algorithm^29^ and transcripts are called by a combination of direct detection and prediction with short reads, using the reference genome as template. The TAPIS pipeline, does not need Illumina reads, but performs several rounds of mapping and correction of RoIs on the reference genome, with apparently similar error correction efficiency as a short-read based method^30^. Finally, the ToFU PacBio pipeline^31^, obtains auto-clusters of FL and nonFL RoIs and then computes a consensus transcript sequence where errors are significantly reduced. In all cases comparison to the reference gene models serves to call known and novel transcripts.

All PacBio transcriptome papers discover thousands of new transcripts, propose a classification scheme by comparing to a reference annotation and find that the majority of novel transcripts appear in known genes^23,25,28,30,32^. However, details on the number, quality and characteristics of these new calls can vary greatly. Sequencing the transcriptome of hESCs by long reads followed by IDP analysis identified over 2,000 novel transcripts (∼30%) and discovered new genes that were proven to be functional^28^. Tilgner *et al.* found using PacBio sequencing of the GM12878 cell line about 12,000 novel transcripts fully supported by previous splice site annotations or Illumina reads, but did not study novel junctions in detail^25^. For the sorghum transcriptome, 11,342 (40%) novel transcripts were found by PacBio from a total of nearly 1M reads using a filter on splice junction quality (SpliceGrapher^33^), and 6/6 random transcripts were confirmed by PCR. Finally, a maize multi-tissue transcriptome analysis identified over 111,151 transcripts from 3.7M RoIs, most of them novel and tissue-specific^32^. The authors found that between 10% and 20% of the PacBio junctions lacked coverage by Illumina reads and < 1% were non-canonical^32^, but do not report on the number of affected transcripts or validate any. In all these cases, an in-depth characterization of the novel transcripts and junctions that would reveal potential biases and justify analysis choices was missing. We believe that such analysis is important as a great variety of FL and nonFL RoIs typically map at each genome locus and different processing pipelines can result in significantly different final transcript calls. As an example, sequencing the mouse neural transcriptome with PacBio, we obtained ∼ 90,000, 13,000 and 16,000 different transcripts when applying Tapis, IDP or the ToFU pipelines, respectively. Implementing a comprehensive, quality aware analysis of PacBio reads is fundamental at a time when long read transcriptome sequencing is becoming more popular and important conclusions on transcriptome diversity will be drawn from these data.

In this work, we present SQANTI (Structural and Quality Annotation of Novel Transcript Isoforms), a pipeline for the analysis of long-read transcriptomics data that defines up to 47 different descriptors of transcript and junction properties, creates a wide range of summary graphs to aid in the interpretation of the sequencing output and implements a machine learning algorithm to remove artifact transcripts based on these descriptors. We apply SQANTI to the analysis of the mouse neural transcriptome and illustrate its usefulness for characterizing transcript types. The robustness of the method is demonstrated by application to several long-read pre-processing pipelines and datasets, and by extensive validation by RT-PCR. The SQANTI analysis confirms the expression of many novel transcripts but also reveals that an important fraction of long-read transcript sequences is presumably alignment or retrotranscription artifacts that can be removed using SQANTI tools. We provide insights into the biological relevance of these new transcripts by describing the biological processes they belong to, characterizing their CDS and UTR diversity, and showing their relevance to accurate transcriptome quantification. Our work confirms the potential of long-read sequencing for precise characterization of the transcriptome complexity provided appropriate preprocessing steps are applied.

## RESULTS

### Experimental design and transcriptome sequencing

Full-length cDNA from Neural Progenitor Cells (NPCs) and Oligodendrocyte Progenitor Cells (OPCs), two biological replicates each, was obtained and split to prepare Illumina and PacBio sequencing libraries (Figure 1A).PacBio sequencing was performed according to the Iso-Seq protocol to generate around 0.6 M RoIs per sample for a total of 2.2M RoIs. Illumina sequencing resulted in approximately 60 M reads per sample. All PacBio RoIs were joined and processed by the ToFU pipeline^31^ to obtain a total of 16,104 primary transcripts. Alignment of the ToFU transcripts against the mouse reference genome (GMAP^34^, assembly mm10) showed an average percentage of coverage and identity above 99.8%, suggesting that the PacBio nominal high raw read sequencing error is corrected by the ToFU clustering approach, as reported^30^. However, small indels (average size ∼ 1.2 nts) were still detected in 56.2% of the transcripts. These small indels did not affect the overall long read mappability as long reads with and without indels had no significant differences in the GMAP quality of mapping parameter and occurred with no particular sequence context bias (Supplementary Figure 1A), which is in agreement with the random profile of PacBio sequencing errors^35,36^ We first attempted to correct indels with matching Illumina short reads using Proovreads^37^and LSC^29^. Although the number of transcripts with at least one indel decreased to 16%, this was still unsatisfactory for ORF prediction. Instead, transcripts were corrected using the reference genome sequence (Figure 1A). By virtue of this strategy, all indels were removed and we obtained the *corrected PacBio transcriptome*.

**Figure 1.**
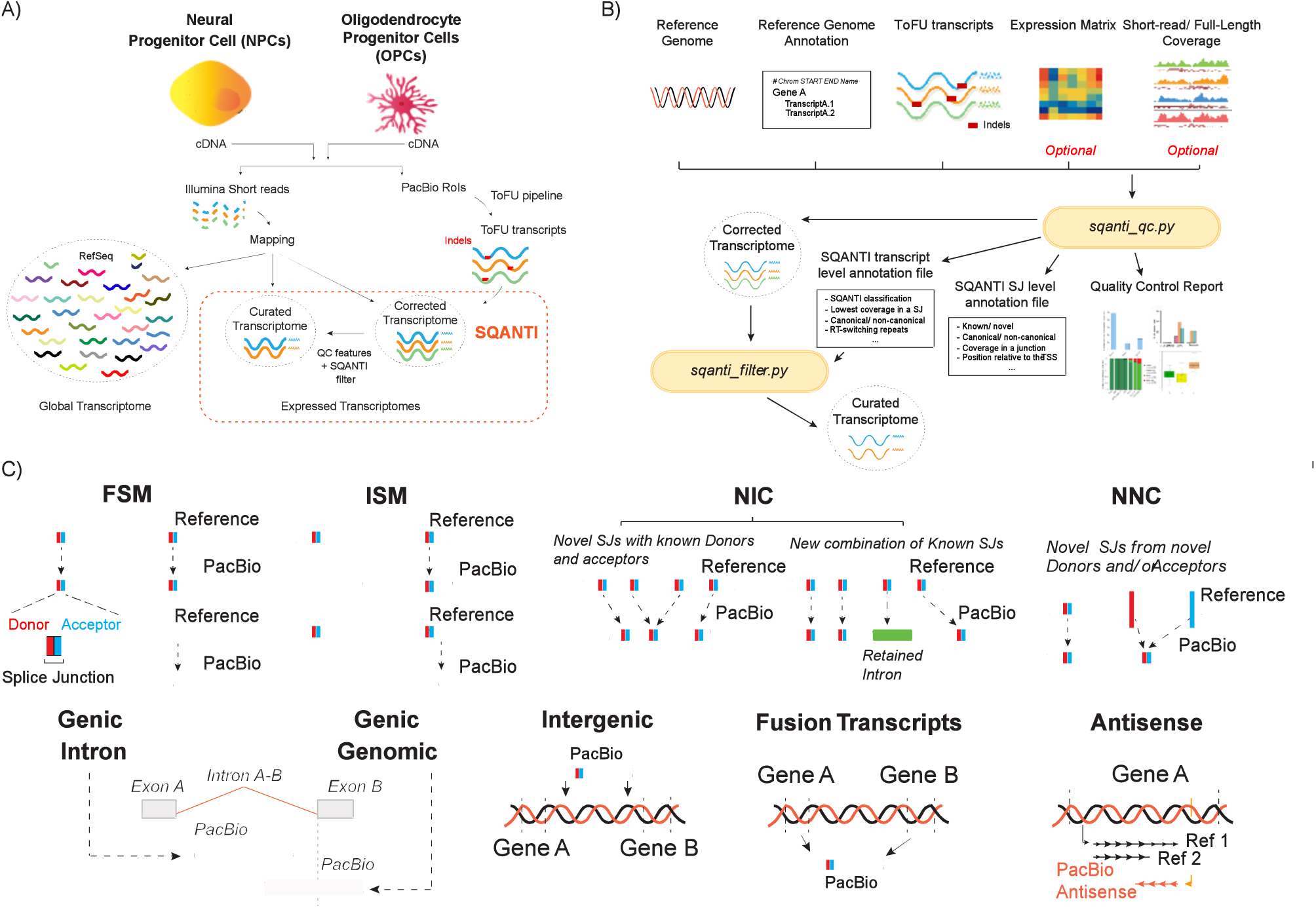
Overview of the experimental model and SQANTI analysis. **A)** Experimental system and data processing pipeline. RNA isolated from Neural Progenitor Cells (NPCs) and Oligodendrocyte Precursor Cells (OPCs) was retrotranscribed separately into cDNA, and sequenced both by long-read PacBio and short-read Illumina technologies. All PacBio RoIs were joined and processed by the ToFU pipeline to obtain consensus transcripts. Residual (indel) errors were eliminated by comparison to the reference genome to generate a *corrected transcriptome* and false transcripts were removed using SQANTI filter to result in a *curated transcriptome*. Illumina short reads were mapped against the RefSeq murine transcriptome annotation, the corrected and the curated PacBio transcriptomes. **B)** SQANTI workflow. Two main functions are part of SQANTI. *sqanti_qc.py* uses as input files a fasta file with transcript sequences, the reference genome in fasta format, a gtf annotation file and optionally Full-Length and short-read expression files. The function returns a reference-corrected transcriptome, transcript-level and junction-level files with structural and quality descriptors and a QC graphical report. *sqanti_filter.py* takes the reference-corrected transcriptome and the transcript-level descriptors file to return a curated transcriptome where artifacts have been removed. **C)** SQANTI classification of transcripts according to their splice junctions and donor and acceptor sites. Splice donors and acceptors are indicated in red and blue respectively. SJ=splice junction, FSM=Full Splice Match, ISM=Incomplete Splice Match, NIC=Novel in Catalog, NNC=Novel Not in Catalog

### Transcript classification based on splice junctions

The SQANTI pipeline was developed for an in-depth characterization of PacBio transcripts. SQANTI takes as input genome and reference annotation information, and returns a reference corrected transcriptome together with a wide set of transcript and junction descriptors which are further analyzed in several summary plots to assess the quality of the data (Figure 1B). Supplementary Tables 1 and 2 describe in detail the set of descriptors computed by SQANTI at the transcript and junction levels, respectively.

A hallmark of the SQANTI analysis is the classification of transcripts based on the comparison of their splice junctions with the provided reference transcriptome to reveal the nature and magnitude of the novelty found by long-read sequencing. For our data we used as reference a non-redundant combination of the RefSeq and Ensembl mouse genome annotations, although other references may be provided by the user. PacBio transcripts matching a reference transcript at all splice junctions are labeled as Full Splice Match (FSM, Figure 1C), while transcripts matching consecutive, but not all, of the splice junctions of the reference transcripts are designated as Incomplete Splice Match (ISM, Figure 1C). Monoexonic transcripts matching a monoexonic reference were included in the FSM category whereas those matching a multiexonic reference were placed in the ISM group (Figure 1C). Furthermore, SQANTI classifies novel transcripts of known genes into two categories: Novel in catalog (NIC) and Novel not in catalog (NNC, Figure 1C). NIC transcripts contain new combinations of already annotated splice junctions or novel splice junctions formed from already annotated donors and acceptors. NNC transcripts use novel donors and/or acceptors. Note that this transcript classification scheme captures the intron-based definition described by^38^, but SQANTI goes one step beyond by describing and sub-classifying the type of novelties introduced by transcripts not matching the splice pattern of annotated references. Novel genes are classified as “Intergenic” transcripts, if lying outside the boundaries of an annotated gene, and as “Genic intron” transcripts if lying entirely within the boundaries of an annotated intron (Figure 1C). In addition, “Genic genomic” category encompasses transcripts with partial exon and intron/intergenic overlap in a known gene (Figure 1C). Finally, SQANTI labels Fusion transcripts (transcript spanning two annotated loci), and Antisense transcripts (polyA containing transcripts overlapping the complementary strand of an annotated transcript, Figure 1C). Our corrected neural PacBio transcriptome contained a total of 16,104 transcripts resulting from the expression of 7,704 different genes. Following the SQANTI classification, transcripts mapping a known reference (FSM, ISM) accounted for 60% of the transcriptome, novel transcripts of known genes (NIC, NNC) made up 35.6% of our sequences. Novel gene transcripts (Intergenic and Genic intron categories) represented about 2.3% of our data while Antisense and Fusion transcripts amounted to 1.1% and 0.3% respectively (Supplementary Figure 1B).

An important advantage of full-length transcript sequencing is that the prediction of ORFs, 5’UTR and 3’UTRs is greatly facilitated. SQANTI implements the GeneMarkS-T^13^ (GMST) algorithm to predict ORFs from transcript sequences which showed highly reliable protein prediction in our data (Supplementary Methods and Supplementary Figure 1C-E). GMST found 11,999 non-redundant ORFs within a total of 14,395 coding transcripts while 1,709 transcripts were predicted to be “ORF-less”. The great majority of FSM, ISM, NIC and NNC transcripts were predicted to have an ORFs (97%, 90%, 87.8% and 92.8%, respectively), while the rest of transcript categories were mostly non coding.

### Descriptive analysis of transcriptome complexity and full-lengthness made easy by SQANTI

A fundamental goal of long-read transcriptome sequencing is to capture the extent of transcriptome complexity and to obtain full-length transcripts. SQANTI includes all basic graphics to readily study these aspects. SQANTI calculates transcript length distribution, reference transcript length, number of supporting FL reads, transcript expression, reference coverage at both 3’ and 5’ ends, number of exons and number of transcripts per gene (Supplementary Table 1). Moreover, analyses are provided with the transcript classification breakdown, which adds an extra layer of understanding on the quality of the sequencing results. For example, we hypothesize that ISM transcripts are a combination of potentially real shorter versions of long reference transcripts as well as partial fragments resulting from incomplete retrotranscription or mRNA decay. We explored the later possibility by analyzing how many ISM transcripts where contained by 95% or more within the UTR3 sequence of their cognate reference transcript and labeled this ISM subclass as UTR3 Fragment transcripts. Indeed, SQANTI analysis shows that PacBio transcripts classified as ISM matched reference transcripts that were longer (Figure 2A) and had more exons (Supplementary Figure 2A) than FSM sequences. Moreover, UTR3 Fragment transcripts matched the longest reference transcripts (Figure 2A) suggesting their enrichment in retrotranscription fragments. All transcript classes had similar median length, except for Genic Intron that was significantly lower (t-test p-value (p) < 1.421 x 10^-15^), while this class and all novel gene categories except fusion transcripts were almost entirely composed by monoexon transcripts (Supplementary Figure 2B). In addition, SQANTI calculates the extent of overlap between sequenced and reference transcript at 3’ and 5’ ends as a proxy to evaluate transcript full-lengthness. This analysis makes sense for FSM transcripts, for which a reference with an identical splice pattern exists. In order to exclude 3’/5’ overlap differences due to alternative polyadenylation or alternative use of TSS we restricted our analysis to matches within the 100 most 5’ or 3’ end nucleotides of the reference. As expected, the majority of our FSM transcripts showed a complete or close to complete 3’ end overlap with the 3’ end of the matched reference transcript: 76% had an exact 3’ end match and 16% were within 20 nts upstream of the annotated 3’ end (Figure 2C). This contrasts with the 35% of FSM transcripts showing a complete overlap with their reference 5’ ends and the 50% falling short by 40 to 100 nts (Figure 2D). This result is in agreement with the strategy used in cDNA library preparation and ToFU analysis parameters that require identification of poly(A) tails to call FL reads, but have less control over completeness at 5’ ends. Interestingly, 851 and 1,361 FSM transcripts had 3’end and 5’end positions further down/upstream of the matched reference transcript, while 1,610 and 1,439 of our FSM sequences, lacked 3’ and 5’ overlap, respectively, of more than 100 nts. These cases might represent alternative polyadenylation/alternative TSS events. Regarding novel genes, only 13.8% of them had splice junctions (Figure 2E) and most (98.2%) expressed just one transcript (Supplementary Figure 2C).

**Figure 2.**
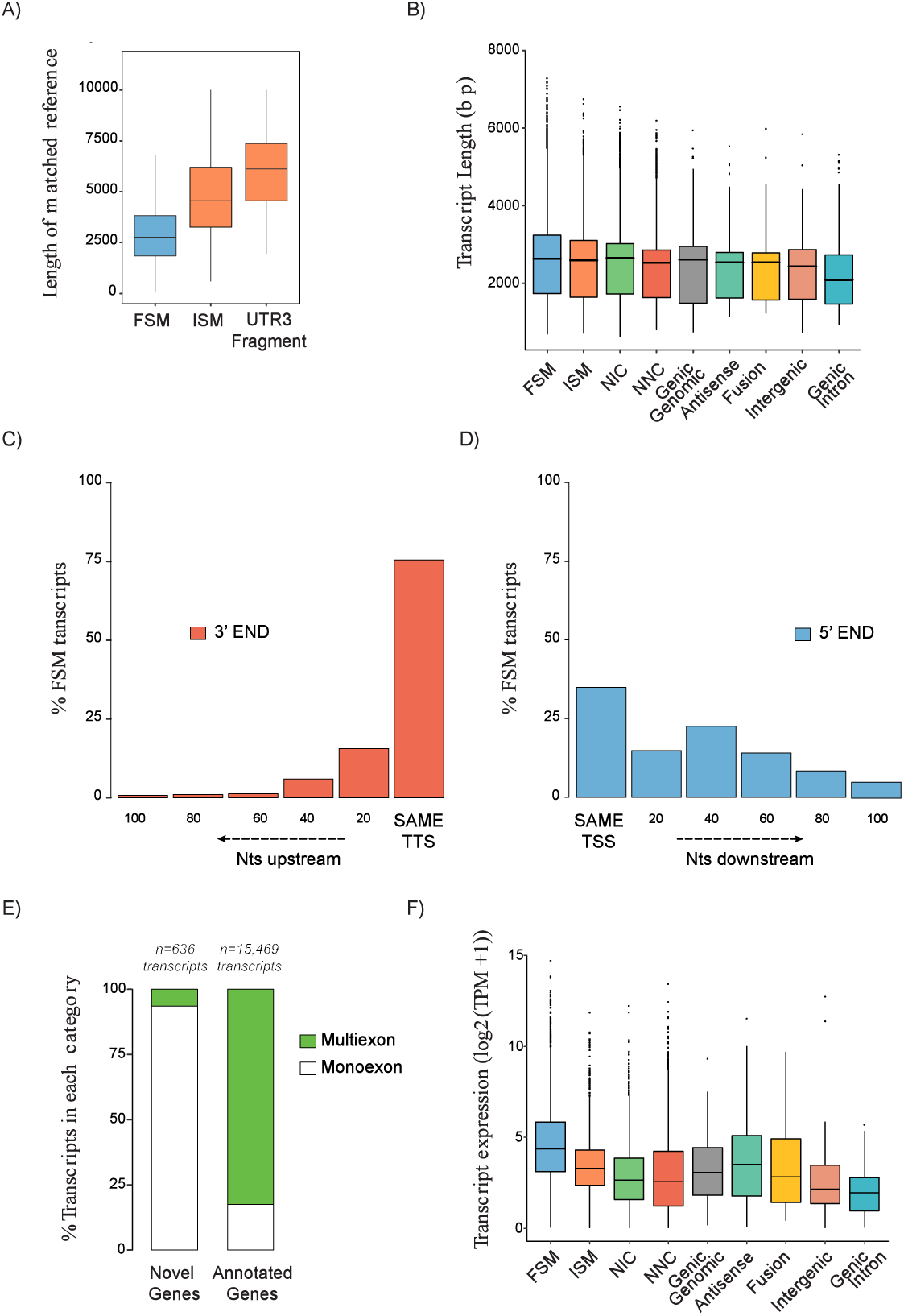
SQANTI characterization of the corrected PacBio transcriptome. **A)** Length of the reference transcripts to which FSM, ISM and UTR3 Fragment PacBio transcripts match. **B)** Length of PacBio transcripts by SQANTI categories. **C)** Overlap at 3’ and **D)** 5’ ends between the FSM transcripts and their respective matched reference transcripts. TTS=Transcription Termination Site, TSS=Transcription Start Site. **E)** Percentage of monoexonic and multiexonic transcripts for transcripts belonging to novel genes and annotated genes. **F)** Transcript expression distribution across SQANTI categories.

Finally, SQANTI descriptive graphs reveal differences between transcript categories at expression features. For example, transcript expression level and number of supporting FL reads were significantly lower in ISM, NIC and NNC transcripts compared to FSM (Figure 2F and Supplementary Figure 2D, t-test p < 2.2 x 10^-16^ for all comparisons) and were significantly lower for novel genes compared to annotated genes (Supplementary Figure 2E and 2F, t-test p < 2.2 x 10^-16^ for both comparisons), which shows that novel transcripts have generally lower expression levels than those already identified in reference databases.

In summary, the descriptive analysis framework provided by SQANTI readily indicates that our neural mouse transcriptome, obtained by PacBio single molecule sequencing, is effective in recovering full-length transcripts and shows an important level of novelty (∼ 40%) with respect to the reference mouse transcriptome both because of novel splicing events and of 3’/5’ end length variation. Transcript diversity is more important than the presence of novel genes, which represents only a small fraction of the expressed mRNAs. However, novel transcripts tend to be less expressed than annotated transcripts indicating that, generally, less novelty is to be expected for major transcripts.

### Evaluation of transcripts according to their splice junctions

In order to better understand the sources of novel transcripts SQANTI further analyzes their splice junctions. Splice junctions can be divided into canonical and non-canonical according to the two pairs of dinucleotides present at the beginning and at the end of the introns encompassed by the junctions. The combination of GT at the beginning and AG at the end of the intron is found in 98.9% of all the introns in the human genome^39^. We considered GT-AG as well as GC-AG and AT-AC as canonical splicing (altogether found in more than 99.9% of all the human intron 39,40), and all the other possible combinations as non-canonical splicing. SQANTI also allows users to provide their own set of canonical junctions. At the same time, SQANTI subdivides splice junctions between known, if they were present in the reference, and novel, if they were not.

In our mouse neural data, the ratio of canonical versus non-canonical splicing events fitted the expected genome proportions when looking at known splice junctions: out of 141,332 known splice junctions, 99.9% were canonical and 0.1% (185) were non-canonical. However, novel splice junctions showed a much different distribution: out of 3,837 novel splice junctions, 69% were canonical and 31% (1,188) were non-canonical. When analyzed across the different SQANTI categories, non-canonical splicing was maintained at low rates in FSM (0.1%) and ISM (0.25%) transcripts, what was expected as both are formed purely by known splicing events (Figure 3A). In NIC transcripts, where novel combinations of known splice junctions or novel splice junctions deriving from annotated donors or acceptors are present, the percentage of non-canonical splicing was 0.15% (Figure 3A). In all cases, these non-canonical junctions were already known in the reference and consequently all novel junctions found in this transcript category were canonical. However, in NNC transcripts, characterized by the introduction of alternative donors and/or acceptors, we found 1,155 novel non-canonical junctions, which represented 4.5% of total. Moreover, Genic Genomic, Intergenic, Genic Intron and Antisense transcripts, despite rarely being multiexonic, showed relatively high percentages of non-canonical splice junctions with 2.32%, 7.28%, 21.57% and 32.65% respectively (Figure 3A). This unusual high level of non-canonical junctions suggests that experimental artifacts might be accumulating in these categories. Furthermore, when the percentage of transcripts showing at least one non-canonical splice junction was considered, the proportion of NNC affected compared to NIC transcripts became more evident, 41.5% vs. 1.47%, respectively (Fisher’s Exact Test (FET) p < 2.2 x 10^-16^), strongly indicating that this category of transcripts needed deeper inspection.

**Figure 3.**
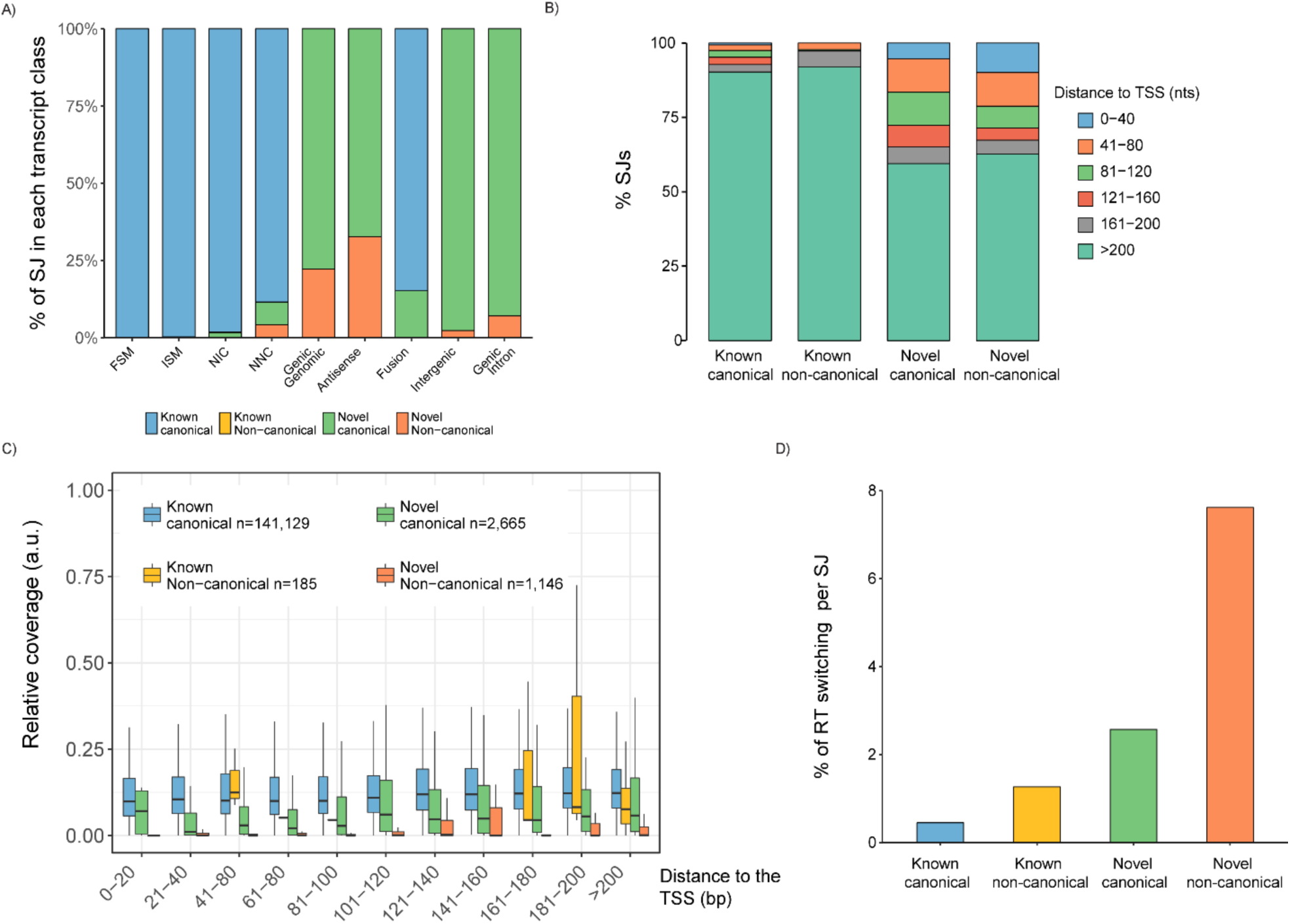
Splice junction’s characterization in the corrected PacBio transcriptome. **A)** Distribution of Splice Junction (SJ) types across SQANTI categories. NNC, Genic genomic, Antisense, Intergenic and Genic intron are enriched in non-canonical SJs. n=76,757 SJ for FSM, n=13,802 for ISM, n=27,368 for NIC, n=26,509 for NNC, n=51 for Genic Genomic, n=49 for Antisense, n=494 for Fusion, n=86 for Intergenic and n=55 for Genic Intron. **B)** Distribution of the SJs according to their distance to the Transcription Start Site (TSS). **C)** Relative coverage by short-reads of SJs as a function of their class and distance to the TSS. a.u. = arbitrary units. **D)** Detection of RT switching direct repetitions by SQANTI algorithm across SJ types.

SQANTI also investigates the position of novel junctions with respect to transcript 5’ ends. We found, that although novel junctions could appear at any position in novel transcripts, there was a higher concentration of occurrences towards 5’ ends which is not observed for known - whether canonical or not - junctions (Figure 3B, FET p < 2.2 x10^-16^). This could either be the consequence of unannotated variability at 5’ ends or higher accumulation of errors due to lower sequence support. The ToFU pipeline is more permissive with clustering conditions at transcript ends (E. Tseng, personal communication), which accounts for a higher probability of errors at these areas.

Coverage by Illumina has been used to support novel junctions called by PacBio^28^. However, Illumina reads are not always equally distributed along the transcript length and are often less abundant towards 5’ ends, providing less support for junction validation. SQANTI integrates short-read coverage data and study the support level of known and novel junctions as a function of their distance to the 5’ end of the PacBio transcript. We found that, as suspected, splice junction support by short reads decreased towards the 5’ end of the transcripts, but was significantly higher for known junctions (Figure 3C, Wilcoxon Rank Sum test (WRS) p < 2.2 x10^-16^). Novel canonical junctions were in general less frequently covered but still significantly more supported than novel non-canonical junctions, which had hardly any supporting reads if located within the first 120 nts of the transcript 5’ end (Figure 3C, WRS p < 2.2 x10-16).

Another possible explanation for non-canonical splicing is Reverse Transcriptase template switching (RT switching). RT switching is an intrinsic property of RTs that allows them to jump within or across template positions without terminating DNA synthesis. Secondary structures in the RNA template have been shown to enhance the RT switching activity^40,41^ and cause gaps during cDNA synthesis. These gaps are interpreted as splicing events, which, due to their non-splicing origin, are enriched for non-canonical junctions^40,41^. A hallmark of RT switching is the presence of a direct repeat between the upstream mRNA boundary of the non-canonical intron and the intron region adjacent to the downstream exon boundary^40^. SQANTI incorporates an algorithm to locate these direct repeats, which confirmed the enrichment of RT switching in novel splice junctions (Figure 3D, FET p < 2.2 x10-16) and in NNC compared to NIC transcripts (7.24% versus 1.98%, FET p < 2.2 x10-16). Described RT switching events affect minor isoforms of genes co-expressed with a major isoform that serves as the template for the intramolecular switching^40^. Accordingly, we found that NNC transcripts are enriched for being minor transcripts of highly expressed genes (Supplementary Figure 2G and 2H).

Finally, SQANTI evaluates possible off priming of the oligo(dT) in A-rich regions of the mRNA template. Annealing of the oligo(dT) primer used in the first strand synthesis of the cDNA to non polyA tail Adenine stretches present in not yet discarded intron-lariats or (pre)-messenger RNAs results in false cDNA molecules^42,43^. SQANTI investigates these events by calculating the % of Adenines (A) within a window of nucleotides downstream of the genetic coordinates corresponding to transcripts 3’ ends. A-rich genomic DNA regions downstream the TTS were concentrated in the relatively minor SQANTI categories (Supplementary Figure 2I), and were enriched in non-coding and monoexonic transcripts (WRS p < 2.2 x10^-16^ for all tests, Supplementary Figure 2J). A total of 601 transcripts were found to be intra-priming candidates, which affected specially the Antisense and Genic Intron categories (∼50% and ∼ 30% of their transcripts were flagged). Remarkably, Incomplete Splice Match (ISM) transcripts that were shortened versions of the reference transcripts by the 3’ end (labeled as 3’ end fragment transcripts) and monoexon NIC transcripts with intron retention events, were also significantly enriched in intra-priming candidates, (WRS p < 2.2 x10^-16^ for all tests, Supplementary Figure 2I).

Altogether, the SQANTI framework analyses suggests that a fraction of the novel transcripts found by ToFU pipeline could originate by technical artifacts at the step of cDNA library construction or by less confident sequencing data at the 5’ ends of transcripts.

### PCR validation of PacBio transcripts

To shed light into the correct detection of transcripts by ToFU analysis we performed RT-PCR amplifications for a total of 67 mRNAs encompassing different SQANTI categories: 23 FSM (3 with non-canonical splice sites), 12 NIC, 30 NNC canonical (11 of them containing at least one non-canonical splice junction) and 3 fusion transcripts (Supplementary Figure 3). Importantly, we performed RT-PCRs both on the ClonTech oligo(dT) enriched full-length cDNAs used for PacBio sequencing and, for positive NIC/NNC/Fusion and 4 FSM transcripts, on new cDNA retrotranscribed at 42 °C and 50 °C using random hexamers rather than oligo(dT). The rationale behind this approach was to test whether novel transcripts could have been spuriously generated by RT switching-like mechanisms at the retrotranscription step of the PacBio protocol. Since higher temperature and/or the use of random hexamers would complicate the formation of secondary structures in the RNA template, retrotranscription artifacts would be less favored in these conditions.

We validated by RT-PCR 23/23 of the FSM, including the 3 cases with non-canonical junctions, (Figure 4A1) highlighting the high level of confidence supporting these transcripts. Novel transcripts showed lower validation rates: 8/12 NIC, 1/3 Fusion and 6/30 NNC, highlighting the low detection rate within NNC category (Figure 4A2). Importantly, 9 of these non-validated NNC transcripts were amplified by oligo(dT) PCR but lost when random hexamers and higher temperatures were used (Figure 4A3), suggesting the occurrence of possible retrotranscription artifacts. Table 1 summarizes the results of the PCR validation experiment. Details can be found in Supplementary Table 3. These results indicated the need of applying an additional filtering strategy to remove artifact transcripts from the ToFU transcriptome output.

**Figure 4.**
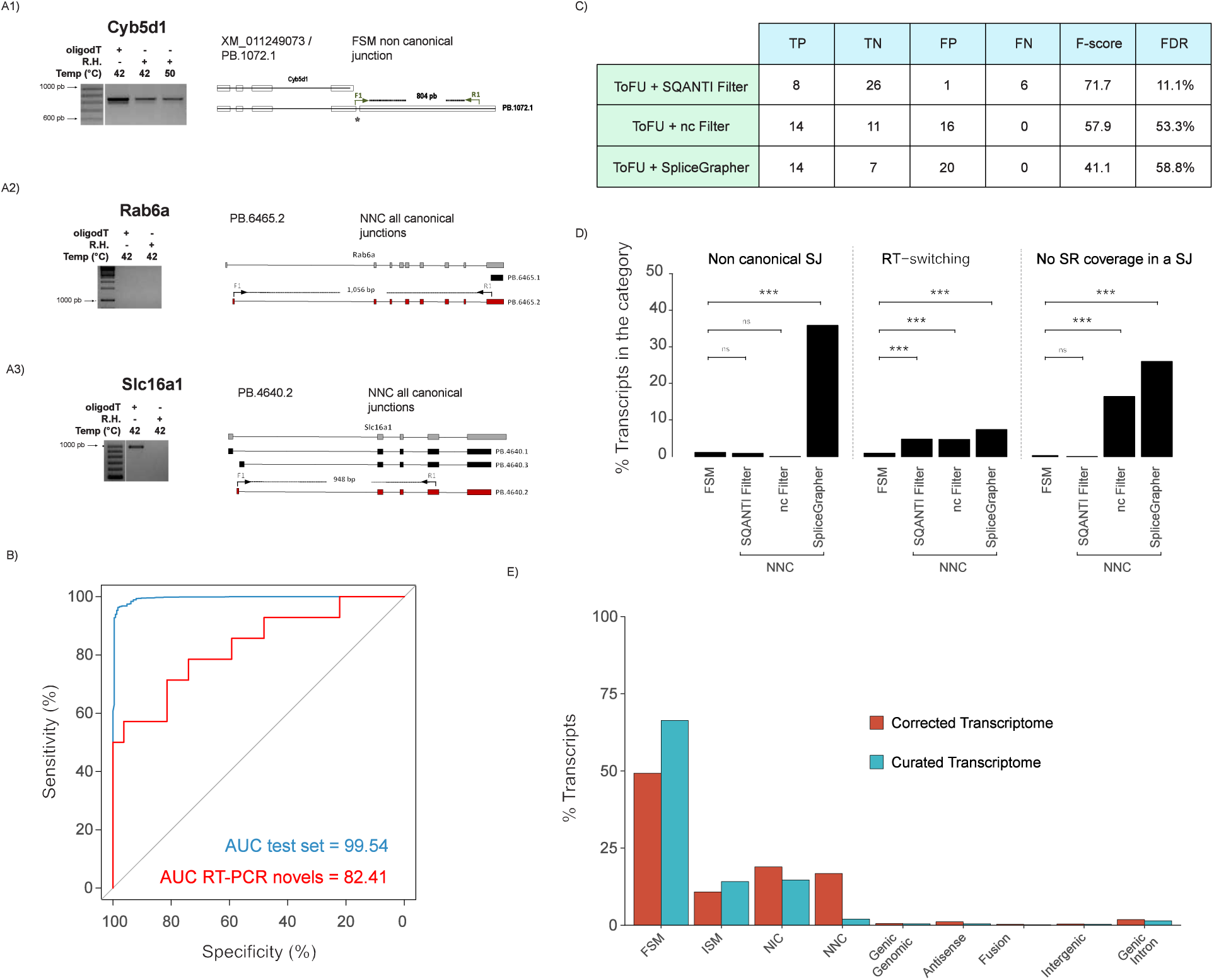
SQANTI filter performance on mouse dataset. **A)** Representative examples of RT-PCR validation experiments. Three PCR conditions were assessed: oligo(dT) template at 42 °C and Random hexamers (RH) template at 42 °C and at 50 °C. **A1)** Example of a FSM transcript with a non-canonical SJ successfully amplified at each PCR condition, **A2)** Example of a NNC transcript with a non-canonical SJ that failed to be amplified in the oligo(dT) condition, **A3)** Example of NNC transcript with non-canonical SJ amplified at oligo(dT) but not at RH conditions. **B)** ROC curves of the SQANTI ML filter for the test set (blue line) and for the set of novel isoforms assayed by RT-PCRs (red line). **C)** Summary of the performances of the SQANTI filter, non-canonical filter and SpliceGrapher filter for the set of novel isoforms assayed by RT-PCR. nc filter = non-canonical filter, TP = True Positive, TN = True Negative, FP = False Positive, FN= False Negative, FDR = False Discovery Rate. **D)** Comparison of quality features in the FSM and NNC categories after SQANTI, nc and SpliceGrapher filters. Statistical differences by Fisher’s Exact Tests (FET), * p < 0.05, ** p <0.01, *** p < 0.001, ns = not significant. **D)** Composition of SQANTI transcript categories in the mouse before and after SQANTI filter.

**Table 1.** Summary RT-PCR validation. nc: transcript with non-canonical junctions.

| Transcript Type | oligo(dT) |  |  | Random Hexamers |  |  | Overall Validation |
| --- | --- | --- | --- | --- | --- | --- | --- |
|  | Positive | Negative | Total | Positive | Negative | Total |  |
| FSM | 23 (3 nc) | 0 | 23 | 4 (3 nc) | 0 | 4 | 100% |
| NIC | 10 | 1 | 11 | 8 | 2 | 10 | 67% |
| NNC | 15 (3 nc) | 15 (8 nc) | 30 | 6 | 9 (3 nc) | 15 | 20% |
| Fusion | 1 | 2 | 3 | 1 | 0 | 1 | 33% |

### Using SQANTI features to build a quality control filter for ToFU artifacts

Previous work applied different criteria to discard artifacts from transcriptome sequencing, including support by short reads^29^, removal of transcripts with non-canonical splicing^38^ or filtering based on sequence features^33^. However, we found that these approaches do not fully capture the complexity of the data. For example, a few known and some junctions in NIC transcripts lack Illumina coverage (148 out of 67,610, and 20 out of 437 respectively), while many of the novel non-canonical junctions did have supporting Illumina reads (543 out of 597). We found that additional features such as RT switching direct repeats and low expression values accumulated in NNC transcripts, but were not exclusive to them. Moreover, our RT-PCR analysis revealed an important number of transcripts (16) having a full set of canonical junctions but failing validation.

We hypothesized that the set of SQANTI descriptors should be informative of transcript confidence and could be used to define a composite filter to efficiently remove artifact transcripts, and decided to train a machine learning (ML) classifier based on these features. To obtain a generally applicable filter we trained our classifier with a “best guess” of true and artifact transcripts within the genome corrected ToFU output. Hence, we defined as positive the Full Splice Match transcripts (FSM, n=7,774) and as negative the Novel Not in Catalogue transcripts with at least one non-canonical junction (NNC-NC, n=1,110), but trained the classifier without providing this structural information (Methods). We used Random Forest^44^ with a 80/20 training/test set split, random down-sampling for class balance and 10x cross-validation, and called predicted transcripts those with a probability for positive classification higher than 0.75. As a note, the RT-PCR instances mentioned in the previous section were excluded from the classifier build. We obtained an Area Under the Curve (AUC) for the ROC curve of the test set of 99.54% (Figure 4B, blue line) and of 82.41% for the set of NIC/NNC transcripts assayed by RT-PCR (Figure 4B, read line). This result indicates that our classifier built on SQANTI descriptors faithfully captures differences between our ground truth set of positive and negative transcripts, which can be efficiently applied to discriminate true transcripts from artifacts within the set of long-read defined novel sequences. Figure 4C shows the performance comparison for the RT-PCR data of the SQANTI classifier against two previous approaches used to remove artifacts, namely the “non-canonical splice junction” filter and SpliceGrapher. Data indicate that our approach has higher F1 score (71.7 versus 57.9 and 41.1 respectively), and lower FDR (11% versus 53.3% and 58.8% respectively) rate than alternative methods. These notable FDR differences are mostly due to a high rate of false canonical junction transcripts that are not discarded by the comparing approaches. Moreover, SQANTI was the only filtering strategy that succeeded in lowering both the *non-canonical SJ* and the *no short read coverage* features in NNC transcripts to levels similar to the high confidence FSM category (Figure 4D).

Features selected by the SQANTI classifier are shown in order of importance in Supplementary Figure 4. The feature ranked first in order of importance (Bite) flags transcripts that skip consecutive exons and have donor/acceptor sites inside a known exon, which we interpret as an indication of novel splice junctions caused by secondary RNA structures. Interestingly, five out of the eight top variables were associated with transcript expression, namely lowest Illumina coverage at junction, minimum sample coverage, number of FL reads, expression of the gene, expression of the transcript and ratio of transcript versus gene expression, suggesting that expression patterns are within the most definitive characteristics to call *bona fide* novel transcripts.

Based on this result, SQANTI incorporates a function for transcriptome curation that applies our ML strategy to the user-provided data by learning classifier parameters on SQANTI descriptor values of each dataset, and filtering accordingly. To compile with additional QC evaluations, the SQANTI filter also includes an option to discard transcripts flagged as intra-priming candidates. Applied to our data, the combination of the SQANTI ML and intra-priming filters removed 4,134 novel transcripts (2,462 NNC, 1,281 NIC, 32 Genic genomic, 36 Fusion, 116 Antisense, 25 Intergenic, 129 Genic Intron and 53 ISM). In our final *curated transcriptome* the adjusted percentages of each category were: 66.3% FSM, 14.1% ISM, 15.7 % NIC, 2% NNC, 0.5% Genic genomic, 0.5% Antisense, 0.2% Fusion transcripts, 0.3% Intergenic and 1.4% Genic Intron (Figure 4E). The transcript category where our filter has the strongest impact is NNC that went from 14% to 2%, while FSM increased consequently from 49% to 66% in the curated transcriptome (Figure 4E). In our final dataset 9,626 transcripts (80.4%) are in the known categories, 2,344 (19.6%) are novel transcripts of which 207 (1.7%) fall within novel genes. These transcripts were the product of 7,167 genes and resulted in 9,269 different ORFs.

### Generalization of the SQANTI approach

To assess the general utility of SQANTI, we applied our approach to alternative analysis pipelines and datasets. We processed our raw mouse PacBio reads with the IDP and TAPIS pipelines and analyzed resulting transcriptomes with SQANTI (Supplementary Figure 5A-B). IDP, which relies heavily on a high quality reference annotation and on short reads correction, returned a total of 13,525 transcripts, the great majority belonging to the FSM category (96%). Only 509 transcripts were novel in this approach (358 NIC, 158 NNC), yet they still showed significant enrichments in RT switching and no short read coverage in a junction (Supplementary Figure 5A). Interestingly, IDP fails to return any of the sixteen novel transcripts validated by PCR, suggesting that this method is highly restrictive for novel isoform calling. On the contrary TAPIS, that as ToFU works without short-read data, returned a significantly larger set of transcripts (91,428) with an overwhelming majority of them belonging to the NNC class (66%), which were strongly enriched in bad quality features (Supplementary Figure 5B).

We next evaluated our analysis pipeline in additional datasets, namely the maize ear^32^ and the human MCF-7 cells^45^, both publicly available. Transcriptome composition in these datasets was substantially similar to what we observed for our mouse transcriptome with a significant number of novel transcripts in known genes, that were enriched in bad quality features (Supplementary Figure 5C-D). We applied the SQANTI filtering approach to these datasets by training our ML classifier in each case with their sets of FSM and NNC-NC transcripts and using default values for removing of intra-priming events. As with the mouse data, we obtained high ROC values in the test sets (99.3 for maize ear and 99.7 for MCF-7) and succeeded in removing a considerable amount of low quality novel transcripts while controlling their enrichment in bad quality features (Supplementary Figure 5 C-D). Additionally we analyzed variable importance of SQANTI descriptors for the ML classifier in these datasets with respect to the mouse data. Interestingly, although we observed an overall agreement in top ranked classification features (i.e. the top three variables were shared among datasets), we also found some noticeable differences (Supplementary Figure 4). For example, the number of FL reads was not a highly ranked feature for the maize ear data, probably due to the lower sequencing depth of this dataset, and was absent in for the MCF-7 dataset, as the value was not available. Still, in both cases high classification performance was achieved. We conclude that our SQANTI filtering approach based in the composite utilization of quality descriptors is a robust but versatile approach for effectively removing artifacts in long read transcriptome datasets that can be applied to a wide range of organisms.

Altogether, this section shows that the SQANTI quality control framework is a very useful tool to reveal the structural composition of transcriptomes obtained from long read sequencing and to compare quality across preprocessing pipelines and experiments. We show that our choice of ToFU read clustering plus SQANTI filtering for transcriptome curation is a good trade-off between discovery and high quality of novel transcript calls, which can be efficiently been applied to different PacBio long read datasets provided that a reference genome and short read data are available.

### Functional insights from novel and alternative transcripts

Most of our novel transcripts belong to existing genes. To further understand the biological relevance of these new calls we analyzed the cellular processes where they participate. Interestingly, genes with novel isoforms were enriched in metabolic processes, regulation of neurogenesis, oligodendroglial lineage, behavior and regulation of potassium ion transport (Figure 5A) suggesting that unannotated isoform diversity impacts both fundamental energy utilization and specific neural biology pathways relevant to our cell type, both key processes for neural differentiation^46–48^. The availability of a full-length corrected and curated transcriptome allows us to predict with high confidence ORFs, annotate 3’ and 5’ UTR and study to which extent alternative splicing modifies coding and non-coding regions of transcripts, Approximately, 36% of the genes expressed in our system were multi-isoform genes. Of these, 1836 genes expressed the Principal Isoform^49^ (PI) of the gene and in 592 cases of these (32%), the PI was expressed in multiple transcripts with variable UTR regions, while for 1,429 genes (79%), an alternative predicted ORF was expressed. In contrast, non-PI transcripts were much less variable at UTRs, with only 9% of them showing multiple 3’ or 5’ UTR variants, and about 27% of the novel transcripts extended existing TSSs or TTSs. This result suggests that in our system, multi-isoform expression mostly resulted in a change in the predicted protein and to a lesser extent in the alternative processing of UTRs. However, alternative ORFs rarely were expressed as more than one transcript, suggesting further transcriptional regulation of these alternative forms might not be required to modulate their functionality.

**Figure 5.**
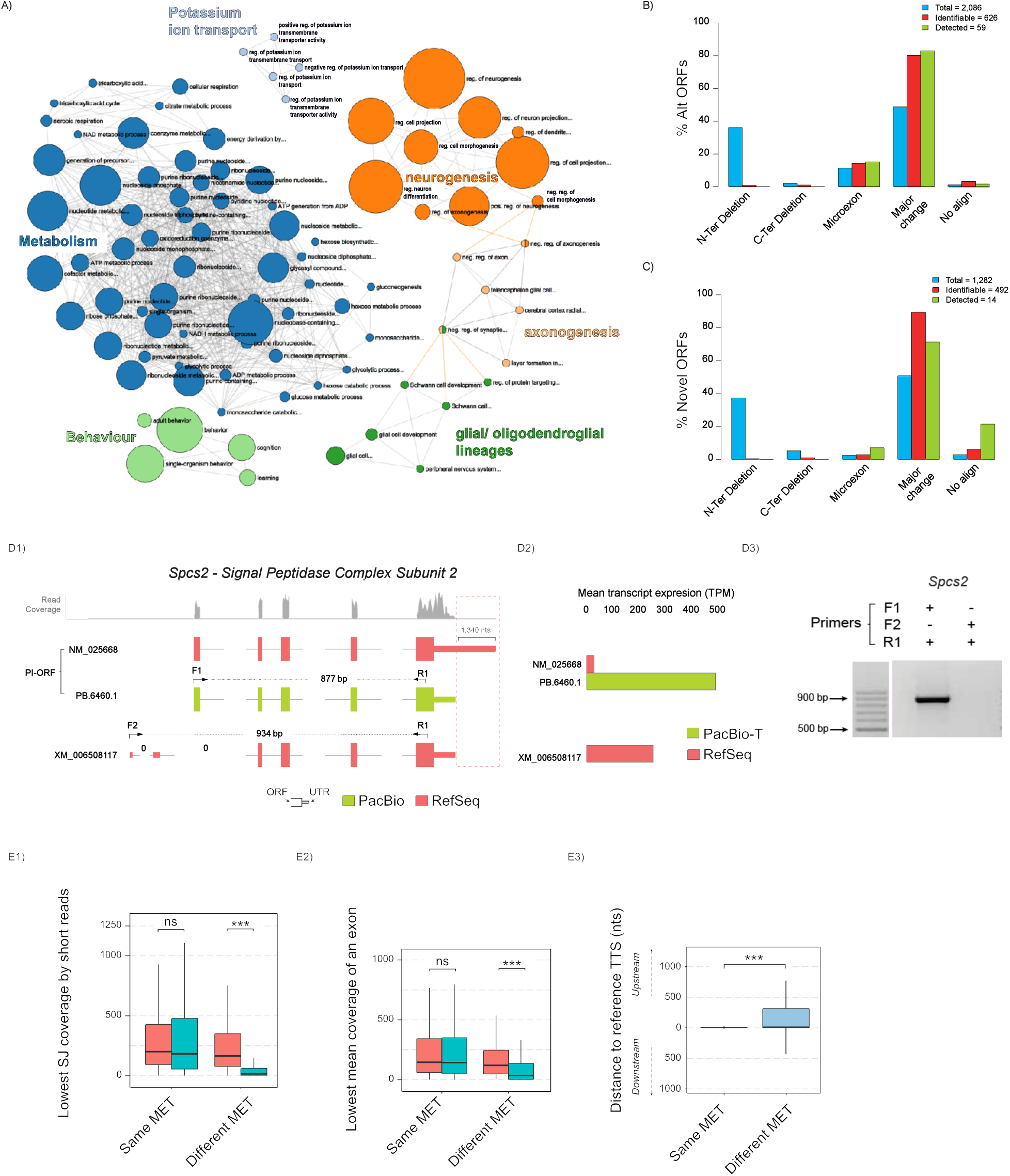
Functional diversity associated to genes with novel isoforms, variability of 3’ UTR in PI-ORFs isoforms and comparative analysis of protein differences between PI and non PI-ORFs. **A)** Gene Ontology enrichment analysis for genes expressing novel isoforms. Analysis of the type of protein changes introduced by **B)** Alternative ORFs and **C)** Novel ORFs with respect to the PI-ORF of the gene. Blue: ORFs computationally predicted in the curated transcriptome; red: ORFs predicted to be identifiable by unique peptides; green: ORFs detected in proteomics databases with at least two Peptide Spectrum Matches (PSMs). **D)** Example of 3’UTR variability in a PI-ORF that leads to a quantification error. **D1)** Transcripts associated to gene Spcs2 gene according to PacBio sequencing (green) and by RSEM quantification using RefSeq (red). The profile of mapping short reads at the Spcs2 locus is shown in grey. Position of transcript-specific primers are indicated by arrows. Differences at the TTS are highlighted by a red dashed box, 0 indicates splice junctions lacking any short-read support. **D2)** Short-reads based average transcript expression levels of *Spcs2* isoforms using either RefSeq or PacBio-T references. **D3)** Validation of Spcs2 isoforms expression by RT-PCR: PB.6460.1/ NM_025668 but not XM_006508117 was amplified. **E)** Analysis of the Most Expressed Transcript (MET) in genes with MET differences between PacBio-T and RefSeq quantifications. Kruskal-Wallis Test, (K-W), *** p < 0.001, ns= not significant. **E1)** Lowest SJ coverage by short-reads in METs. **E2)** Lowest mean exon coverage by short-reads in METs. **E3)** Distance between the Transcription Termination Sites (TTS) of the METs and their FSM references. Same MET means both PacBio-T and RefSeq select the same MET, Different MET means RefSeq selects a MET that is not manually curated and PacBio-T selects a MET that is manually curated.

### Peptide support of novel and alternative transcripts

As most of the novel transcripts were predicted to have ORFs that contained novel amino acid stretches when compared to PIs, we sought to investigate whether peptide data present in public proteomics databases could support these findings. In order to do this we first created a non-redundant ORF database of public mouse proteins and the predicted proteins in our neural data, and classified each protein as Principal Isoform ORF (PI-ORFs, n=4,579) if annotated as such in the Principal Isoform predictor APPRIS^49^, Alternative ORF (Alt-ORF, n=2,127), if present in Ensembl or RefSeq but not being PI, and Novel ORF (Novel-ORF, n=1,194), if the protein would be coded by NIC or NNC transcripts present only in our mouse PacBio data. For each predicted protein, we performed an *in-silico* trypsin digestion and selected unique peptides that would unequivocally identify each ORF. We analyzed theoretical peptides for those genes identified in our mouse transcriptome that had more than one isoform annotated in Ensembl (v80). The percentage of ORFs predicted to be identifiable by unique peptides was highest for the PI-ORFs (56.3% or 2,577), followed by the Novel-ORFs (42.6% or 509) and was lowest for Alt-ORFs (30.1%, or 641). The majority of Novel-ORFs and Alt-ORFs were predicted to have only one unique peptide, while this was only the case for 14.2% of the PI-ORFs (Supplementary Figure 6A). Conversely, most PI-ORFs were predicted to contain 6 or more discriminating peptides and this was true for only 7% of Alt-ORFs and 9.8% of Novel-ORFs. This higher rate of unique peptides in PI-ORFs was expected as the mouse genome contains a significant number of genes in which alternative isoforms have only partial sequences and the APPRIS PI is often the longest ORF in a gene. Consequently, proteins deemed as PI are expected to be easier to detect by protein digestion approaches than alternative isoforms.

We then screened public databases for the presence of unique peptides associated to our set of ORFs. Two separate approaches were conducted: a *Neural tissue* approach, comprising one proteomics study of mouse neural tissue and another study of the mouse neural secretome, and an *All tissue* approach comprising peptides from 36 proteomics studies carried out on a variety of murine tissues but excluding the two ones used in the first approach. Overall, we detected at least one unique peptide for 77.9% of the PI-ORFs predicted to be identifiable, while this percentage went down to 20.56% and 8% for Alt-ORFs and Novel-ORFs, respectively. Most Alt- and Novel-ORFs had single unique peptide matches, while most PI-ORFs were found with multiple peptides (Supplementary Figure 6B). In part this is to be expected; the success of detection was significantly lower when the ORF was predicted to have only one unique theoretical peptide, and this was the case for the majority of Alt-ORFs and Novel-ORFs (Supplementary Figure 6C). Interestingly the agreement between the two proteomics screening approaches was much stronger for those proteins detected with two or more peptides (Supplementary Figure 6B). When ORFs were identified by a single peptide, the peptide was almost always present in just one of the two studies. Note that ORF detection by single peptide matches, similarly to transcript detection by single read counts, falls into the area of unreliable protein identification and therefore false discovery in these cases is not controlled^50^. This result confirms that the lower number of discriminating peptides in Alt and Novel ORFs versus their PI ORF counterparts impairs their detection by proteogenomics approaches, but other factors are also contributing, as Alt/Novel ORFs had lower unique peptide detection rates across all unique peptide ranges (Supplementary Figure 6C).

To understand if expression levels were playing a role, we evaluated the number of studies (PSM counts) supporting each ORF to find that on average Alt- and Novel-ORFs had 5-6 supporting studies (median=2) per detected unique peptide, while this number was nearly 10 for PI-ORFs (median=4.5), which is in agreement with the notion that PI-ORFs are ubiquitously expressed across tissues^49^. Interestingly, we found that PI-ORFs detected by unique peptides in less than 5 proteomics studies had a significantly lower expression in our system than those found in more than 10 projects, and had similar expression levels as the transcripts coding for Alt- and Novel-ORFs (Supplementary Figure 6D). Altogether, these results indicate that direct detection in public proteomics databases of predicted coding products of novel and alternative transcripts is hampered by their lower expression pattern and an overall lower identifiability by unique peptides.

Finally, we evaluated the types of protein differences between alternative and principal isoforms for which peptide support was conclusive (minimum of 2 PSM counts per ORF, n=59 Alt-ORFs and n=14 Novel-ORFs), and compared them to the composition of our predicted transcriptomes. While our set of curated transcripts predicted that most alternative and novel ORFs distributed between N-terminal truncations, microexons (indels/substitutions up to 9 amino-acids, aas) and major changes (indels/substitutions of more than 9 aas with or without N-Ter/C-Ter truncations), the proteogenomics analysis failed to reveal most of these N-terminal differences and mostly found the major changes both for Alt- and Novel-ORFs (Figure 5B-C), which is in agreement with a detection approach that relies on positive detection of unique peptides. Microexons were found mostly in Alt-ORFs (Figure 5B) while Novel-ORFs with no overlap to their PIs were found in the proteomics databases more than expected (Figure 5C), however this finding is supported by just a few ORFs and hence cannot be conclusive. Interestingly, although there was more than a 10-fold difference between the number of identifiable ORFs and those consistently identified in our proteomics screenings, there was a general agreement between the relative abundance of each type of protein differences among the two ORF sets, which suggests that the ORFs confidently identified by unique peptide matches could represent the actual diversity range of the alternative proteome.

### Novel transcripts have a major impact on accurate transcriptome quantification by short reads

Previous studies have shown that the utilization of a reduced, expressed transcriptome as reference for short-read mapping instead of the total reference dramatically impacts transcriptome quantification^51,52^ and improves replicability of expression level estimates^28^. We sought to investigate how the new transcripts impact quantification by short-reads. As one important aspect of transcript-resolved analysis is the identification of the transcript that captures most of the expression in each gene (Most Expressed Transcript, MET), we concentrated our study in the comparison of METs when using the total RefSeq (∼160,000 transcripts) or the curated PacBio transcriptome (11,970 transcripts, aka PacBio-T) as reference for short-read mapping. For 3,976 genes the MET was identical in PacBio-T and RefSeq, meaning that the PacBio-T MET was a Full Splice Match of the RefSeq MET. Interestingly, this was not the case for 1,433 genes, and in 996 of them the PacBio-T MET was a different FSM transcript, therefore present in RefSeq. For example, the Signal Peptidase Complex Subunit 2 gene (*Spcs2*) was expressed as one transcript in our PacBio neural transcriptome (PB.6460.1) and had two transcripts in RefSeq quantification (NM_025668 and XM_006508117) (Figure 5D.1). PB.6460.1 is a FSM transcript of NM_025668 and both codify for the PI-ORF of the gene but the 3’ exon of PB.6460.1 is smaller, resulting in a 3’UTR shorter by 1,340 nucleotides, (Figure 5D1, red dashed box). This shorter 3’ exon is actually the annotated exon of the RefSeq transcript, XM_006508117, which also uses two alternative 5’ exons (Figure 5D1). XM_006508117 was the MET in the RefSeq quantification while NM_025668 was estimated as poorly expressed (Figure 5D2). Upon RT-PCR amplification with transcript discriminating primers we confirmed the PacBio-T and not the RefSeq based quantification scheme (Figure 5D3). When inspecting read coverage at this locus we observed that neither the unique 5’ junctions of XM_006508117 nor the extra exonic sequence at the 3’exon of NM_025668 were covered by Illumina short reads, while the short-read pattern nicely fits the PacBio transcript model. We speculate that this variability at the 3’UTRs creates a conflict when resolving transcript quantification in the RefSeq gene model that was decided in favor of transcript XM_006508117 by RSEM^53,54^, as this transcript has a more consistent 3’ end coverage. In summary, the transcript quantification error of the *Spcs2* gene when using a reference transcriptome as mapping template was due to a discrepancy in the 3’end annotation between the reference and the actual expressed transcripts. Similar disagreement patterns were observed for two additional genes, *Dhrs7b* and *Bdkrb2* with similar outcomes in terms of MET selection (Supplementary Figure 6 E and F). To estimate how general this pattern was, for all the MET discrepant genes we investigated the RefSeq curation status. Interestingly, the majority of the discrepant genes (47.2%, n=470 genes) corresponded to situations where the PacBio-T MET was a FSM of a manually curated RefSeq transcript and the RefSeq MET was not manually curated, as in the *Spcs2* gene. Furthermore, in these cases the RefSeq-based MET had significantly worse lowest splice junction coverage and lowest mean exon coverage than the MET called by the PacBio-T quantification (Figures 5E1 and 5E2). Similarly to *Spcs2*, we found that for these 470 genes the differences in the length at the 3’ end between the MET selected at PacBio-T quantification and their matched RefSeq transcripts were significantly higher than in genes where both quantifications selected equivalent METs (Figure 5E3). Moreover, these differences were also observed for transcripts codifying for the PI-ORF of the genes, indicating that the extensive variability in the 3’ ends that is not annotated in a global reference such as RefSeq is not restricted to secondary/alternative transcripts. These results demonstrate the relevance of using a full-length reference transcriptome updated with novel expressed transcripts for correct quantification estimates.

## DISCUSSION

### SQANTI as a critical tool to analyze whole transcriptome quality

Long read sequencing technologies, such as the PacBio platforms as well as Illumina’s Moleculo and Oxford Nanopore, have brought novel excitement into the challenge of describing the complexity of the transcriptome of higher eukaryotes by providing new means for sequencing full-length transcript models. While early papers concentrated on demonstrating the dramatic enrichment in full-length transcripts achieved by long reads^23,55^, there is an increasing number of publications that describe thousands of new transcripts discovered by this technology. Accordingly, we found that, when sequencing the mouse neural transcriptome using PacBio a large number of novel transcripts could be detected. However, close inspection of these new transcripts revealed signs of potential errors that required a thorough and systematic analysis of these sequences before making any new transcript calls. This motivated the development of SQANTI, a new software for the structural and quality analysis of transcripts obtained by long-read sequencing.

The three basic aspects of the SQANTI QC pipeline are i) the classification of transcripts according to the comparison of their junctions to a reference annotation in order to dissect the origin of transcript diversity, ii) the computation of a wide range of descriptors to chart transcript characteristics and iii) the generation of graphs from descriptors data, frequently with a transcript-type break-down, to facilitate interpretation of the sequencing output and reveal potential biases in the novel sequences. Using this analysis framework we were able to show that, at least in our mouse experiment, novel transcripts - especially those in the NNC category - are typically poorly expressed transcripts of known genes and that novel junctions accumulate at the 5’ end of transcripts, have lower coverage by Illumina reads, and are enriched in non-canonical splicing and direct repeats typical of RT-switching. However, none of these features are exclusive of any of the novel transcripts categories, which invites the question on “how best to remove transcript artifacts”. This has been solved in the past by either eliminating all novel transcripts with at least one junction not supported by short-reads^23^, by systematically discarding transcripts with non-canonical splicing^28^ or by developing models to estimate the likelihood of a certain splicing event^30^. In our case, we performed an extensive PCR validation of transcripts belonging to different known and novel types. Surprisingly, we found a significant number of transcripts, both with canonical and non-canonical junctions, that did have complete junction support by Illumina and were amplified by RT-PCR of the sequenced cDNA library, but failed to be validated when PCR conditions were adjusted to avoid secondary RNA structures. We concluded that these might be cases of retrotranscription artifacts, which would have escaped a filtering solely based on short-read support. This result may suggest that a revision of library preparation protocols is needed, which goes beyond the scope of this study. As an alternative, we were able to combine our set of SQANTI descriptors with a machine learning strategy to build a filter that discards poor quality transcripts with better performance than the methods indicated above. The SQANTI filter is data-adaptive and can we showed that it can be successfully applied to other long read transcriptomics datasets. Note that SQANTI is designed to leverage genome annotation data to characterize and filter the long read transcriptome. In case no genome is available or the assembly is low quality, reference-guided correction of transcript sequences will be compromised and hence accurate translation into ORFs. If, additionally, the gene content annotation is poor this will impact SQANTI transcript classification, that will be enriched in novel isoforms and genes. In these conditions it might be difficult to define robust FSM positive and NNC-NC negative training sets for the SQANTI classifier. The first because of the low number of known transcripts and the second because of poor correction of Pacbio sequences. Sub-sampling experiments showed that 150-200 training set transcripts would be sufficient to obtain comparable performance to that in Figure 4B (not shown), indicating that the SQANTI filter can be used confidently even when reduced training sets are available. Furthermore, the SQANTI set of quality descriptors will be extremely useful in these cases, as they will provide a comprehensive characterization of the quality of the transcript calls in situations where little additional data is available.

### Novel insights in transcriptome complexity from single molecule full-length transcriptome sequencing

The fundamental advantage of single molecule long reads technologies over short reads is their direct detection of full-length isoform diversity and of novel transcripts. The availability of a curated full-length transcriptome dataset of our mouse neural tissue allowed us to explore these aspects confidentially. We found that genes with novel transcripts are enriched in metabolic processes and specific neural functions related to neurogenesis and oligodendroglial lineage. This is remarkable because both the narrow control of metabolic programming and the expression of genes involved in cell identity are key players of differentiation courses^56^, and the finding that most novel transcripts concentrate in these categories suggests that an important untapped transcript/regulatory diversity present here could be revealed by long read sequencing technologies. We find interesting to note that most of the transcript diversity is concentrated in the appearance of novel ORFs but also an important fraction of the alternative transcripts are UTR variations of the Principal Isoform of the gene. However, alternative isoforms are rarely also expressed with variable UTRs and novel transcripts infrequently extend annotated TSSs and TTSs. This suggests that gene expression regulation by alternative transcripts either controls the expressed protein or the transcript stability but the interaction of the two might not be as critical. We also show how high variability at transcript ends is a source of quantification errors that can be alleviated when an expressed full-length reference transcriptome is used. Our data suggests that unannotated alternative polyadenylation events are frequent in mammalian genomes, which in turn induce incorrect quantification estimates. Full-length sequencing of the expressed transcriptome readily identifies this 3’end diversity to provide the correct templates for transcript quantification. On the other hand, variability at the 5’ end is still an issue for full-length transcriptome sequencing as biological variability cannot be unequivocally differentiated from technical artifacts in cDNA library preparation protocols. The SMARTer protocol typically used in PacBio sequencing may not always capture the full extension of the 5’ ends due to transcript degradation or incomplete retrotranscription. This may account for the lack of 5’ end coverage observed in FSM and ISM transcripts. Interestingly, trapping of the 5’ CAP prior to the synthesis of the secondary cDNA strand has been shown to increase the overlap of the 5’ end without seriously compromising the yield of long reads^57^ and in future may represent the preferred form of library preparation to study 5’ end diversity.

Finally, we investigated whether the transcriptome diversity found by long read sequencing was mirrored by proteogenomics data. We concluded that the low expression and identifiability by single peptides of Alt and Novel ORFs hampered their detection by proteomics. Detection of alternative protein isoforms has proved to be difficult, and while some authors claim that limited detection in proteomics databases indicates low translational and stability rates^58,59^, other studies identify a significant proportion of alternative exons associated to ribosomes as evidence of active translation^60,61^. While it is not the scope of this work to resolve these issues, we turned our attention to the analysis of protein differences for those cases of confident peptide detection. Interestingly, we found that the distribution of the type of protein differences in the non PI-ORFs with respect to the main isoforms is similar to the predictions based on the PacBio sequencing data, except for N-terminal truncations that are at a disadvantage in a unique peptide detection approach. Most of detected alternative ORFs showed major protein changes compared to the PI-ORF of their respective genes, which could potentially have an impact on functionality of the alternative protein. While a detailed analysis of these functional differences requires further computational and experimental approaches, the results presented in this paper indicate that long read technologies, provided adequate quality control is applied, are effective tools for describing the isoform-resolved transcriptome and can aid in the study of the biological significance of alternative splicing.

## MATERIAL AND METHODS

### Differentiation of NPCs and OPCs from neonatal mice

Neonatal c57/BL6 mice (4 days old) were sacrificed and Neural Precursors cells (NPCs) were isolated from the subventricular zone. Neurospheres were obtained by culturing the progenitors in media supplemented with EGF and bFGF and Oligodendrocyte Precursor Cells (OPCs) were derived from them by adding ATRA (All Trans Retinoic Acid) as described in the Supplementary Methods section.

### RNA extraction, full-length cDNA library preparation and sequencing

Total RNA isolation from cultured cells (two biological replicas per cell type) was done with Nucleospin RNA kit (Macherey-Nagel) obtaining RINs (RNA Integrity Number) between 10 and 9.7 for all samples. The synthesis of full-length cDNA was performed with SMARTer PCR cDNA Synthesis kit (ClonTech, version 040114, CA, USA) following PacBio recommendations. The cDNA synthesis protocol used 1 µg of total RNA, 42 °C for retrotranscription and 13 PCR amplification cycles to control for over-amplification of small fragments. For each sample, we performed two first-strand cDNA synthesis reactions and nine PCR reactions using 10 µl of first strand cDNA (diluted 1:5 in TE-buffer) to obtain around 14-16 µg full-length cDNA per sample. Each sample was submitted to the ICBR sequencing facility (University of Florida) for PacBio sequencing (P4-C2 chemistry). Three cDNA fractions were obtained with BluePippin and sequenced at the RSII Instruments using 2 SMRT cells for the 1-2 kb fraction, and 3 SMRT cells for 2-3 kb and 3-6 kb fractions, to a total of 8 SMRT cells per sample. Additionally, the same samples were sequenced with the Illumina Nextseq instrument using Nextera tagmentation and 2x50 paired end sequencing. Sequencing data has been submitted to the SRA under Submission number SUB2459157 (PacBio reads) and SUB2466432 (Illumina reads) and will be released upon publication of this manuscript.

### Transcriptome generation and quantification

Sequenced PacBio subreads were pooled together and ToFU software was used to obtain non-redundant transcripts. Default parameters were set to obtain Read of Insert (RoI), Full-length classification of RoIs and ICE (Iterative Clustering for Error Correction) steps. Quiver option was turned on to improve consensus accuracy of previously generated ICE clusters by using non Full Length read information. Generated HQ polished isoforms (>99 % accuracy after polishing) were collapsed to eliminate isoform redundancy (5’ different was not considered when collapsing isoforms). This set of 5’ merged non-redundant isoforms was defined as ToFU transcriptome. TAPIS was run with default parameters, except for the maximum intron length used by GMAP (version 2016-05-01), which was set to 200,000. Apart of the reference genome, TAPIS requires the input of a transcriptome annotation file, in this case the RefSeq murine transcriptome. IDP corrects long sequences through the incorporated LSC^29^ module that maps high quality short-reads to Iso-Seq long reads using Bowtie2 (version 2.3.2). The parameters were set to default but for the aligner (GMAP, see command line in Supplementary Methods) and the minimum isoform fraction value to accept a predicted transcript which was set to 5%. Transcript quantification using short-reads was obtained using STAR^62^ as mapper and RSEM^53,54^ as quantification algorithm (parameters available at Supplementary Methods). Expression estimates were obtained as Transcript per million (TPM). Long-read quantification was computed as the number of Full-Length reads of each transcript divided by the total number of FLs of the sample. Most Expressed Transcript (MET) was defined as the transcript of each gene that obtained the highest average TPM value across all the samples. The relative coverage of a splice junction was defined as the sum of all the reads mapped to the junction divided by the sum of the expression of all the transcripts in which it is present.^29^ module that maps high quality short-reads to Iso-Seq long reads using Bowtie2 (version 2.3.2). The parameters were set to default but for the aligner (GMAP, see command line in Supplementary Methods) and the minimum isoform fraction value to accept a predicted transcript which was set to 5%. Transcript quantification using short-reads was obtained using STAR as mapper and RSEM as quantification algorithm (parameters available at Supplementary Methods). Expression estimates were obtained as Transcript per million (TPM). Long-read quantification was computed as the number of Full-Length reads of each transcript divided by the total number of FLs of the sample. Most Expressed Transcript (MET) was defined as the transcript of each gene that obtained the highest average TPM value across all the samples. The relative coverage of a splice junction was defined as the sum of all the reads mapped to the junction divided by the sum of the expression of all the transcripts in which it is present.

### Verification of transcripts by Reverse Transcription PCR (RT-PCR)

PCR amplification of selected transcripts was performed with both the sequenced full-length cDNA and newly synthesized cDNA from the same RNA extractions. For new cDNA reactions, 1 µg of total RNA was used to synthesize the first-strand cDNA using SuperScript III (Life Technologies) primed with random hexamers in a reaction volume of 20 µl, according to the manufacturer’s instructions. Each random hexamer cDNA synthesis reaction was carried out at two temperature conditions: 42 °C and 50 °C. RT-PCR reactions used 1 µl of sequenced full-length cDNA or 2 µl of random hexamers cDNA, together with Biotools DNA Polymerase (1U/ µl) in a reaction volume of 50 µl. Primers were designed to span the predicted splicing event using Primer-BLAST^63^ Supplementary Table 3, http://www.ncbi.nlm.nih.gov/tools/primer-blast). PCR condition were 5 min at 94 °C followed by 35 cycles of 94 °C 30 s, primer-specific annealing temperature for 30 s and 72 °C for 1 min or 1:30 min, depending of predicted product size. PCR amplification was monitored on 1.5 % agarose gel.

### RT switching prediction

SQANTI contains an algorithm that implements the RT switching (RTS) conditions described in Cocquet et al^40^. Namely an exon skipping pattern due to a retrotranscription gap caused by secondary structures in expressed transcripts. The algorithm looks at all the junctions for possible RTS (both canonical and non-canonical junctions) and checks for a direct repeat pattern match at defined sequence locations: the pattern at the end of the splice junction’s 5’ exon must match the pattern at the 3’ end of the splice junction’s intron. There are three parameters that control pattern matching: (1) the minimum number of nts required to match (4 - 10); (2) the number of nts of wiggle allowed from the ideal pattern location (0 - 3); (3) whether allow for a single mismatch, indels or not. SQANTI uses as default parameters: a minimum of 8 bases long repeat sequences, a maximum wiggle of 1 and no mismatches. FSM transcripts with the highest mean expression in each gene are assumed to serve as templates for RTS and are excluded from the analysis.

### ORF prediction and functional annotation

The GMST algorithm^13^ was applied to predict ORFs in PacBio transcripts, setting parameters to only consider the direct strand of the cDNA and AUGs as the initial codon. As GeneMarkS-T allows prediction in incomplete transcripts lack of coverage in the 5’ end caused some truncated ORF starting in codons different from Methionine. In these instances the ORF was shortened by the N-Terminus until the first in frame Methionine was found. GMST was benchmarked as shown in Supplementary methods. GO annotation of novel transcripts was done by Blast2GO^64^ with default parameters and a query-hit overlap requirement of 90% of the hit sequence^65^ and enrichment analysis was performed with the hypergeometric test of the goseq^66^ R package.

### Characterization of Alt-ORF and Novel-ORF with respect to PI-ORFs and UTR/ORF variability

Microexon definition was restricted to novel amino-acid (aa) stretches obtained by in-frame indels or substitutions of no more than 27 nts (9aas) following Irimia et al^67^. ORFs showing exclusively N-Ter deletions or C-Ter deletions were labeled as N-Ter Deletion or C-Ter Deletion ORFs. ORFs showing indels and substitutions greater than 9 aas, combined or not with N-Ter and C-Ter deletions, were labeled as Major Change ORFs. ORFs that could not be aligned against the PI-ORF of their respective genes were deemed as No align ORFs. Two UTRs were considered to be different if they started in different genomic coordinates or if they shared a common start point but had a length difference of more than 30 nucleotides.^67^. ORFs showing exclusively N-Ter deletions or C-Ter deletions were labeled as N-Ter Deletion or C-Ter Deletion ORFs. ORFs showing indels and substitutions greater than 9 aas, combined or not with N-Ter and C-Ter deletions, were labeled as Major Change ORFs. ORFs that could not be aligned against the PI-ORF of their respective genes were deemed as No align ORFs. Two UTRs were considered to be different if they started in different genomic coordinates or if they shared a common start point but had a length difference of more than 30 nucleotides.

### Machine Learning classifier of artifacts based on SQANTI features

A machine learning approach was developed to discriminate artifacts from true novel transcripts utilizing SQANTI features. FSM transcripts were used to define the set of positive transcripts while NNC-non canonical transcripts were taken as negative set. By definition, the labeled sets (FSM and NNC-NC) contain only multi-exonic transcripts, and hence the classifier can only be applied to this type of transcripts. From the total set of SQANTI transcript descriptors, 16 variables defined for both novel and know transcripts sequences were selected (Supplementary Table 1). SQANTI transcript descriptors that relate to reference transcripts, structural category classification and canonical junction status were excluded as either they are irrelevant to the classification or they were used to define the positive and negative transcript sets. Variables with near zero variance or a correlation higher than 0.9 in the labeled sets are removed. The labeled set was divided into a training set (80%) and a test set (20%) and algorithms were run using down-sampling to equilibrate positive and negative sets and 10 times 10 cross validation. Several machine learning methods were tested (Adaboost^68^, CART^69^, Random Forest^44^, SVM^70^, Treebag^71^) on the mouse data that employed 7774 FSM, 1100 NNC-NC transcripts and 14 SQANTI descriptors (*RTS_stage* and *coding* variables were excluded in this dataset due to low variability). Random Forest (RF) was selected as best performing approach (Supplementary Methods) and run using 500 trees. This RF approach was also applied to the PacBio maize ear^32^ and human MCF-7^45^ datasets. For all datasets, the quality of the predictions was assessed by ROC analysis and evaluation of SQANTI quality descriptors on the filtered transcriptome obtained after the application of the classifier to the novel transcripts. For our mouse dataset, SQANTI filter performance was also evaluated on 67 transcripts tested by RT-PCR and computing ROC, F1-score and FDR values. The F-score was calculated as 2*(Specificity*Sensitivity/(Specificity+Sensitivity)). The FDR was calculated as 100*(FP/(TP+FP)). Note that transcripts evaluated by RT-PCR were excluded from training set used to build the classifier.

### SQANTI pipeline

SQANTI is implemented in Python with calls to R for statistical analyses and generation of descriptive plots. The SQANTI program has two major functions: sqanti_qc and sqanti_filter. The sqanti_qc function performs different tasks: (1) corrects transcript sequences based on the provided reference and returns a corrected transcriptome; (2) compares sequenced isoforms with current genome annotation to generate genes models and classify transcripts according to splice junctions (a full-description of structural classification of isoforms can be found in results section); (3) predicts ORFs using GeneMarkS-T; (4) runs our algorithm to predict RT switching; (5) returns a transcript level and junction level descriptive file. These files contain 33 and 20 fields respectively where the three first fields identify the transcript in the reference genome and the remaining fields describe different transcript/junction properties, making a total of 47 SQANTI descriptors (Supplementary Tables 1 and 2). sqanti_filter uses the SQANTI features output to perform filtering of artifacts by two different approaches. The intra-priming filter option removes transcripts with adenine stretches in the genomic position downstream their 3’ end. The machine learning filter learns a Random Forest classifier on the user’s data following the strategy described above. sqanti_filter returns a curated transcriptome where artifact transcripts are removed. For the mouse, maize^32^ and MCF-7^45^ datasets the reference genomes used were mm10, AGPv4 and hg38, respectively. SQANTI is available at <u>https://bitbucket.org/ConesaLab/sqanti</u>.

### Analysis of Peptide Support

We performed an *in silico* analysis of the peptide support of the predicted ORFs of our neural transcriptome when compared to public proteomics databases. A non-redundant database composed of predicted ORFs from our murine transcriptome experiments and all the murine ORFs annotated in Ensembl (v80) was created. These ORFs were subjected *in silico* tryptic digestion (Proteogest, complete digestion). Unique peptides were identified and ORFs with at least one unique peptide of 7 amino acids of more were annotated as *identifiable ORFs*. We then used two different approaches to detect experimental Peptide to Spectrum Matches (PSMs) that match unique peptides from our ORFs. The first approach made use of a pipeline built on Pladipus^72^, a platform that allows for distributed and automated execution of bio-informatics related tasks and performed an all tissue search of mouse proteomic studies (n=36). The pipeline consists of pride-asap, a tool designed to automatically extract optimal search parameters, SearchGUI^73^, a tool that manages the execution of several search engines and PeptideShaker^74^, a tool that allows for the merging of the results produced by the search engines. For this study, X!Tandem^75^, Myrimatch^76^ and MSGF+^77^ algorithms were applied. The input spectra were obtained from 36 murine projects in the PRIDE^78^ database. The second approach was based on the Sequest algorithm^79^ and screened large-scale mouse proteomics experiments of brain tissue^80^ and astrocyte secreted proteins^81^. A more detailed description of these approaches is available in Supplemental Methods.

## ACKNOWLEDGEMENTS

We thank Eric Triplett (University of Florida) for support in sequencing experiments and Elizabeth Tseng (PacBio) for helping in running the ToFU pipeline and critically reading this manuscript. This work has been partially funded by the University of Florida Pre-eminence hires program, the Spanish Ministry of Economy and Competitiveness grant BIO2015-71658-R and Spanish Ministry of Education grant FPU2013/02348.

